# An Integrated 3D bioprinted “disease in a dish” lung cancer model to simultaneously study drug efficacy, toxicity and metabolism

**DOI:** 10.1101/2022.05.02.490270

**Authors:** NP Nandeesha, Madhuri Rotella, Subrahmanyam Vangala, Uday Saxena

## Abstract

Discovery and development of new drugs is a long, expensive and high risk proposition. Millions of dollars spent and decade plus years of time taken to discover a new drug have haunted pharma industry for many years.

In part, the reliance on animal models to make go or no go decisions for selecting drugs for human trials has been a problem because animal biology does not capture human disease in entirety. In recognition of this, the last decade has seen the emergence of more human like tools being developed in the hope of better prediction of human outcomes.

Towards that end we have developed a 3D bioprinted disease in a dish lung cancer model which uses human cells and includes ability to measure drug efficacy, toxicity and metabolism simultaneously. For drug profiling studies in our disease in a dish model we 3D bioprinted intestinal cells, layered below which were liver cells and finally underneath were target lung cancer cells. The idea was to simulate the path taken by an oral drug which encounters the gut, followed by liver and target organs. We demonstrate here that a 3D bioprinted disease model composed of human derived cells is able to concurrently measure in vitro drug efficacy, toxicity and metabolism. Such humanized models will help make early go or no go decisions on the potential of a drug to enter human trials.

## Introduction

3D bioprinting has evolved as superior technology to create human like physiological systems and human organs. The goals of such research are a) long term goal is to create human organs that may be used for transplantation b) current goal is to create 3D systems capable of mimicking human physiology for drug testing.

There are two types of research activities within 3D bioprinting – a bioengineering focus to create the best tools/models and another where the focus is to ask a question and create the 3D model that can enable answering the question. We present here a 3D model which specifically asked the question of simultaneous measurement of drug efficacy, toxicity and metabolism in a lung cancer model.

Drug discovery and development paradigms currently use separate vitro assays and animal models sequentially to profile drug candidates and select lead molecule. For example, to study efficacy usually there are one set of screening assays, to study metabolism there are again separate series of cell free and cell-based assays and finally toxicity is studies in a different set of tests. Compounds emerging from this tedious activity are then tested in animal models which usually do not reflect human disease or metabolism. As a result of using this old-style paradigm the predictive value of models used is poor, time consuming and expensive.

We have designed a 3D bioprinted system consisting of human intestinal cells, liver cells and lung cancer cells (target tissue). There are recently published reports of cancer 3 D printed models but they not incorporate efficacy, toxicity and metabolism in one system. Our hope was that our all-encompassing system may better reflect human efficacy, toxicity and drug metabolism. We also expected that such a “disease in a dish” model may be a better alternative to screen drugs than the series of assays currently used. Finally, the use of human cells may better predict human outcomes rather than animal models currently used.

## Brief Methods

### 3D bioprinting of three layers

Human intestinal cell Colo205 or Caco2, liver cell (Hepg2) and lung cancer cells (A549), (ATCC) were used as representative cell lines and maintained in Dulbecco’s modified or Eagle’s Medium (DMEM, ATCC, USA) supplemented with 15% fetal bovine serum (Thermo Scientific) and 1% Penicillin/Streptomycin. This media was also used for 3D bioprinting. 3D bioprinter from 3D Cultures (Philadelphia, USA) was used. Cells to be bioprinted were loaded in a 10 m l syringe (which serves as a printer cartridge in bioink). Printing pressure and dispenser valve opening times (pulse duration) are critical parameters for the proper printing of various biomaterials, hydrogels, and cells and were optimized for each of the cell types. Typically after printing each layer the plate was put back in incubator for 24-48 hrs and next layer was then printed.

To study the metabolism, efficacy and toxicity of a drug, the drug was added to the 3D bioprinted layers in 12/24 well plates in the media at the top of the well. The media was later collected after defined time, from 0-48 hours, for HPLC analysis using published methods for each drug. The cell layers are examined for cell morphology using a phase contrast microscope.

The viability of cells grown in layers by 3D bioprinting was assessed by addition of MTT and then counting the stained cells under a phase contrast microscope. For every layer at least five representative fields were counted and data analysed by ImageWare. The different layers were counted by adjusting the microscope to focus on a particular layer. In some experiments the cells were then lysed and MTT colour OD was measured using a spectrophotometer.

## Results

### 1. Disease in a dish model development

The cells were 3D printed as shown in **Figure 1 :** target cells lung cancer cells A549 at the bottom, followed by hepg2 liver cells and finally at the very top were the colo205 (or caco2 cells). The cells in each layer were viable up to at least 48 hrs after the assembly of the 3D bioprinted system. So drug properties after addition to the 3D bioprinted system were studied for at least 48 hrs after addition of the drug. During this time a drug can be added once or multiple dosing and can also be added a few hours apart if needed to simulate single dose versus multiple dose regimens.

**Figure 1.**
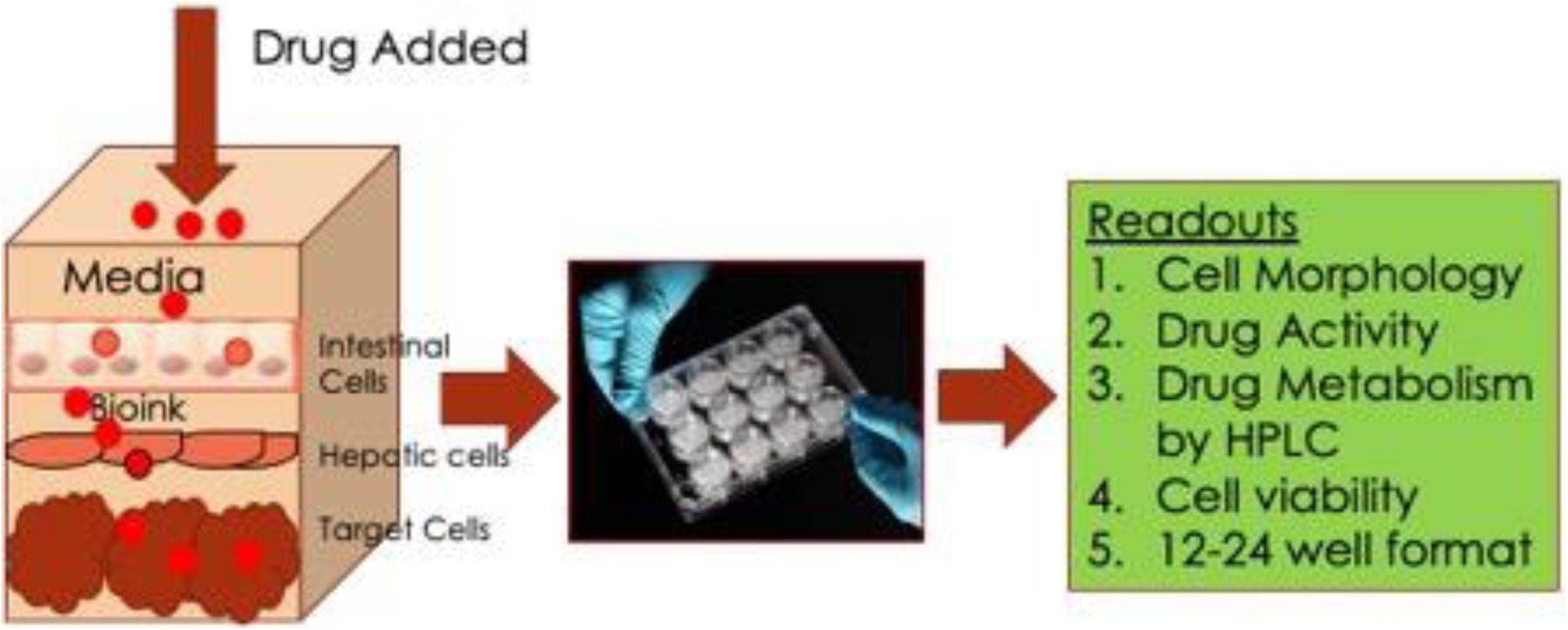
shows the construction of our 3D bioprinted system and end points measured.

Cell morphology by phase contrast microscopy provided status each of the three cell layers. It was found that the cell morphology was in line with what is expected of each layer suggesting that the cells grow well in this 3D dish and are healthy. Cell viability as also assessed using MTT and showed all three cells types to be viable for at least 48 hours after all layers are bioprinted.

### 2. Efficacy scoring of drugs in the 3D system

Since lung cancer A549 cells were used, this system can be used to study drugs that are active against lung cancer. Three groups of drugs for validation were selected - those that are known to be chemotherapeutic, those that have shown some activity and then those that were not expected to have any activity.

The data provided herein shows that the 3D bioprinted system shows more effective activity by cancer chemotherapy drugs but not by other two groups of drugs. The reproducibility of the system for demonstrating drug activity was ascertained by demonstrating that similar degree of inhibition (IC50 around 40uM) was obtained in three separate experiments using cisplatin, which is a known anti-cancer chemotherapeutic **(Figure 2)**.

**Figure 2.**
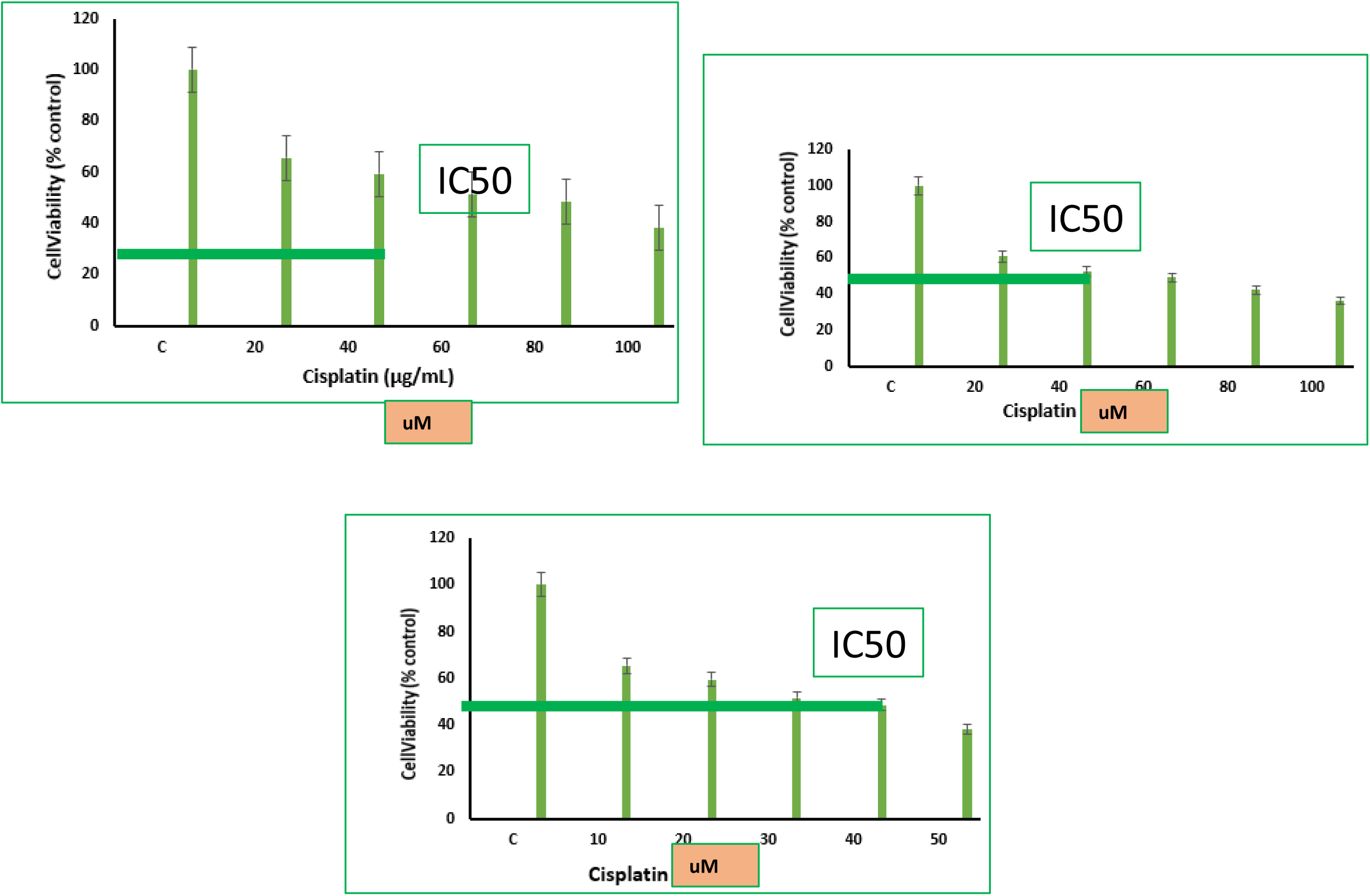
shows IC50 data from three separate experiments in which cisplatin activity was tested and found to be reproducible.

### 3. Efficacy of various drugs against lung cancer A549 cells in the 3D bioprinted system

As shown in **Table 1**, the chemotherapeutic drugs cisplatin and cyclophosphamide were most active as shown by their IC50 (drug concentration needed for 50% of activity). We also profiled other classes of structurally diverse drugs as mentioned above such as pro drug cyclophosphamide, a chemotherapeutic which needs to be converted to an active metabolite by the liver cells.

**Table 1.** shows IC50 values of various drugs tested for anti-cancer activity in 3D system.

| Drug Name | IC<sub>50</sub> | Comments |
| --- | --- | --- |
| Cisplatin | 45µM | Active |
| Cyclophosphamide | 12.5µM | Active |
| Indomethacin | 50 µM | Moderately Active |
| Paracetamol | >100 µm | No consistent activity |
| Metformin | No activity | Negative control |
| Testosterone | No activity | Negative control |

Indomethacin shows moderate activity as has been shown to be active in some clinical studies but there are no reports of its activity in PDX animal models. Interestingly, this drug is found to be active in the present 3D system **(Table 1)** establishing that this 3D model correlates with human data.

Metformin and testosterone which are not effective for lung cancer showed no activity. Overall these data strongly support that the present system is reflective of drug activities against lung cancer and indomethacin activity supports activity similarity to human reports.

### 4. Toxicity Index of various drugs in 3D bioprinted model

Besides testing efficacy, the present 3D printed platform is also used to simultaneously test the general toxicity of the test drug or drug combination.

Toxicity was analysed by looking at a drug’s activity on the target A549 cells **(Figure 3)** versus any effect on the liver hepG2 cells **(Figure4)**. Any killing of the hepG2 cells is considered general toxicity since it is not the target cell. It was then scored for cell death of the target A549 cells versus the liver Hepg2 cells. We calculated the ratio of cell death ratio between heg2 cells: target A549 cells. A higher ratio would indicate more/selective death of the target cancer cells relative to the liver Hepg2 cells. This ratio is termed as Toxicity Index of the drug. As shown in **Table 2**, the chemotherapeutic drugs cisplatin and cyclophosphamide which work by being cytotoxic has a ratio of close to 1, suggesting that they killed the targetA549 cells and the liver HepG2 cells equally. In contrast indomethacin had a ratio of more than 6 suggesting that it was mainly killing the target A549 cells but not the liver HepG2 cells. Paracetamol which is known to be hepatotoxic showed a ratio of 2 suggesting some toxicity toward liver cells. Metformin and testosterone tested as negative controls did not kill either liver Hepg2 or A549 cells. Thus, our system can be used to look at efficacy and toxicity at the same time and establish a Toxicity Index.

**Figure 3.**
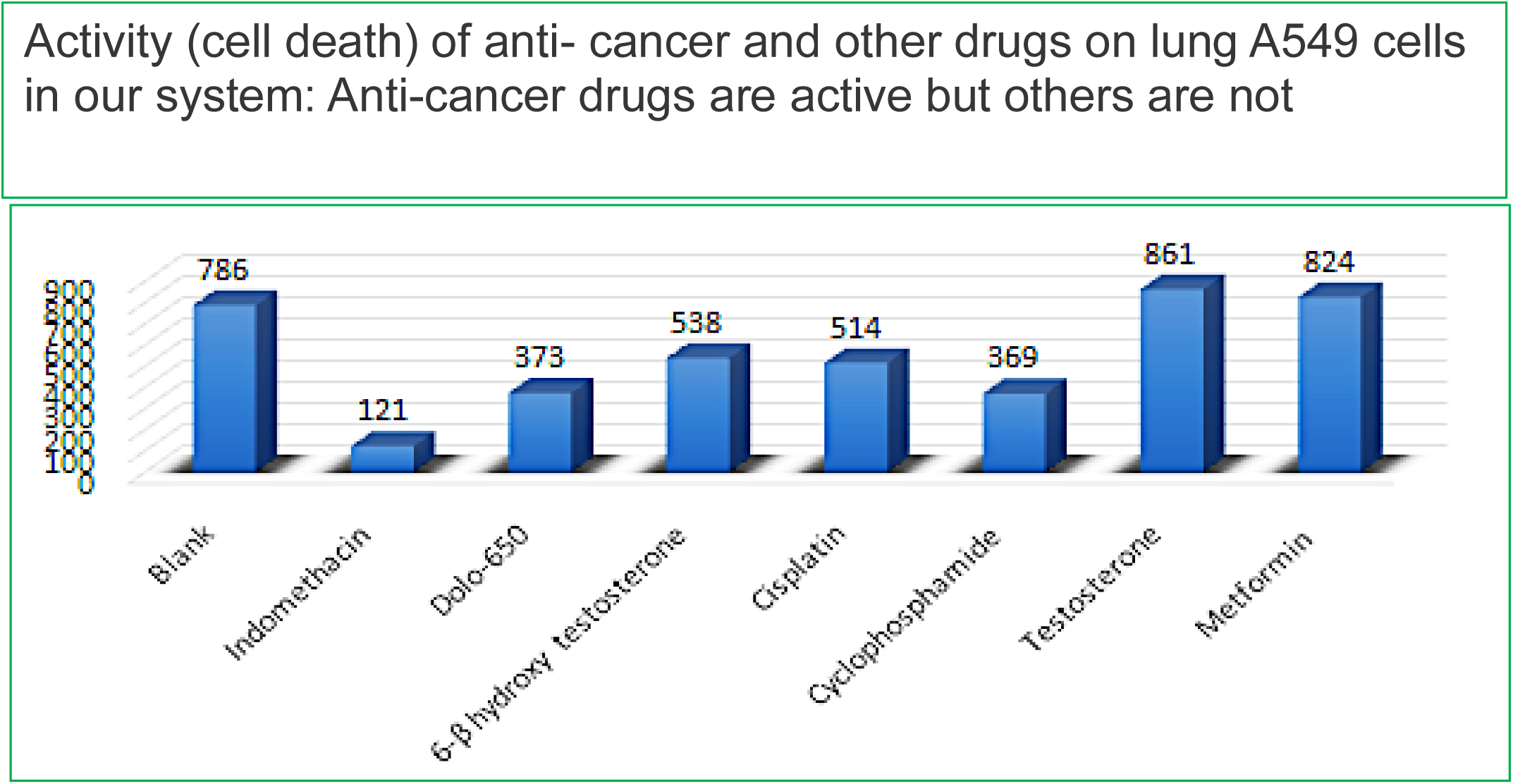
shows the cell counts of lung cancer cells after drug treatment. Lower cell numbers indicate higher cell death. Blank bar represents untreated cells.

**Figure 4.**
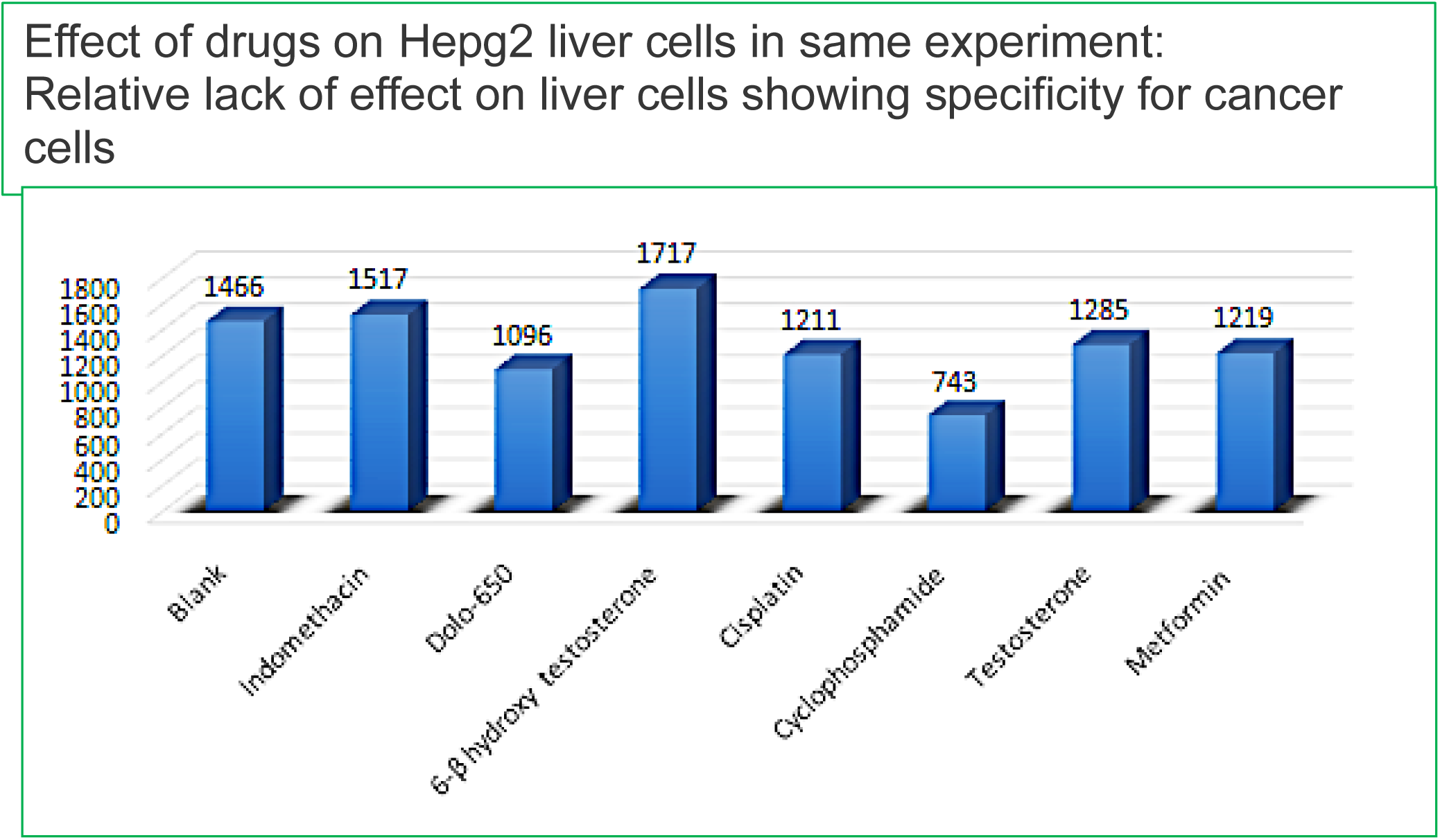
shows liver cell counts after drug treatment. Blank bar represents untreated cells.

**Table 2.**

| Name of drug | Toxicity Index (ratio of cytotoxicity to liver Hepg2 cells: cancer A549 cells) | Comments |
| --- | --- | --- |
| Indomethacin | 6.6 | Indomethacin was selectively toxic to cancer cells but not liver cells |
| Paracetamol | 2.2 | Drug was more toxic to cancer cells but also to liver cells |
| Cisplatin | 1.07 | Similarly toxic to both cell types |
| Cyclophosphamide | 1.08 | Similarly toxic to both cell types |
| Testosterone | No quantifiable death in either cell type | - |
| Metformin | No quantifiable death in either cell type | - |

These data show that using our 3D bioprinted system we can simultaneously look at activity/toxicity of target and other cells

### 5. Analysis of Drug metabolism using 3D bioprinted system versus lung cells alone

One of the advantages of our 3D bioprinted system is the ability to study metabolism of the test drug alongside efficacy and toxicity. The metabolite profile of drugs tested above in the 3D bioprinted system versus just the target cells A549 cells alone was compared using the media to detect metabolites by HPLC.

A s shown in **Figures 5-8**, it was predominantly found that the metabolic activity in 3 D bioprinted system was distinct and superior in terms of the number of metabolite peaks seen as compared to incubating drugs with target cells A549 cells alone.

**Figure 5.**
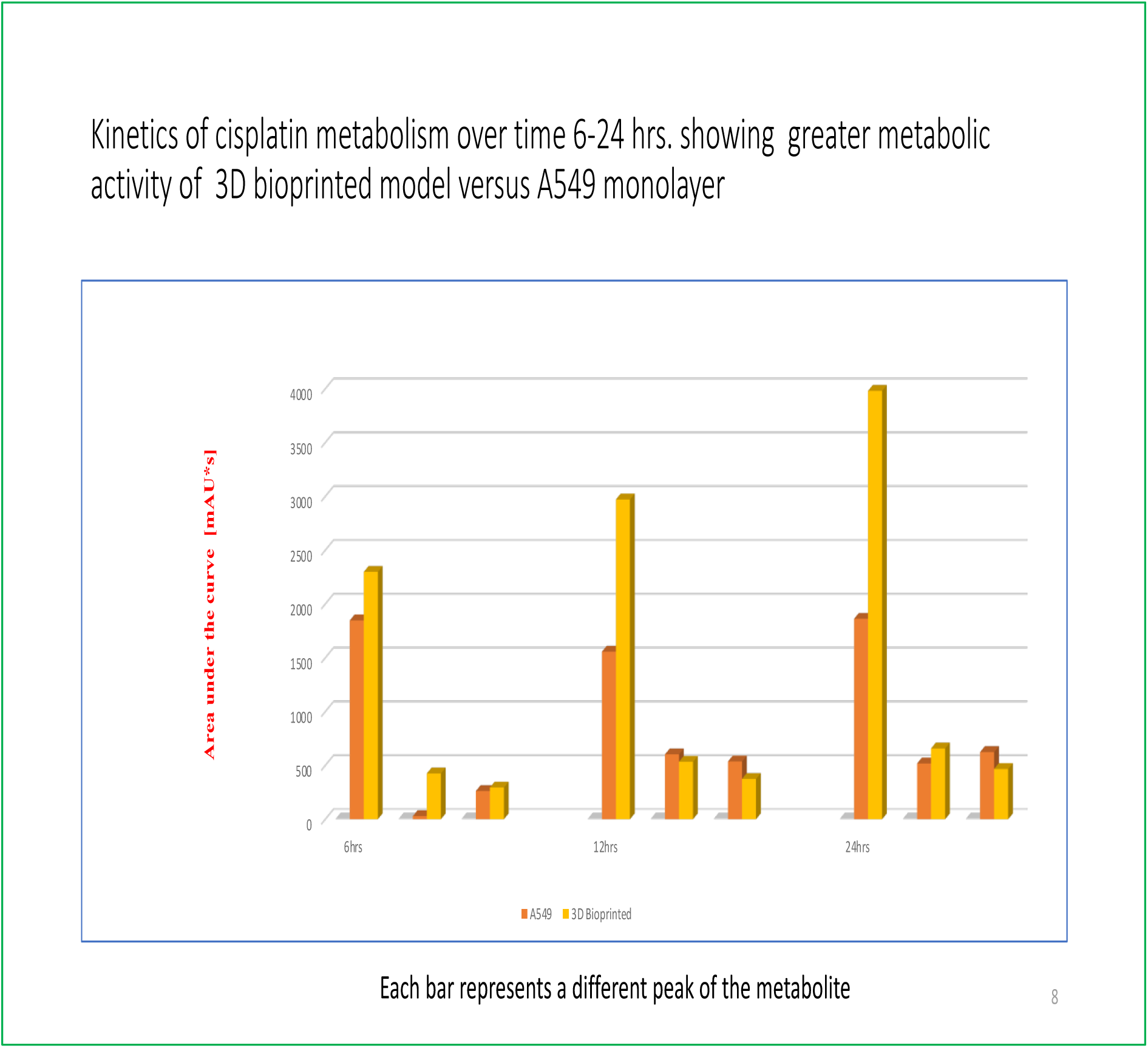

**Figure 6.**
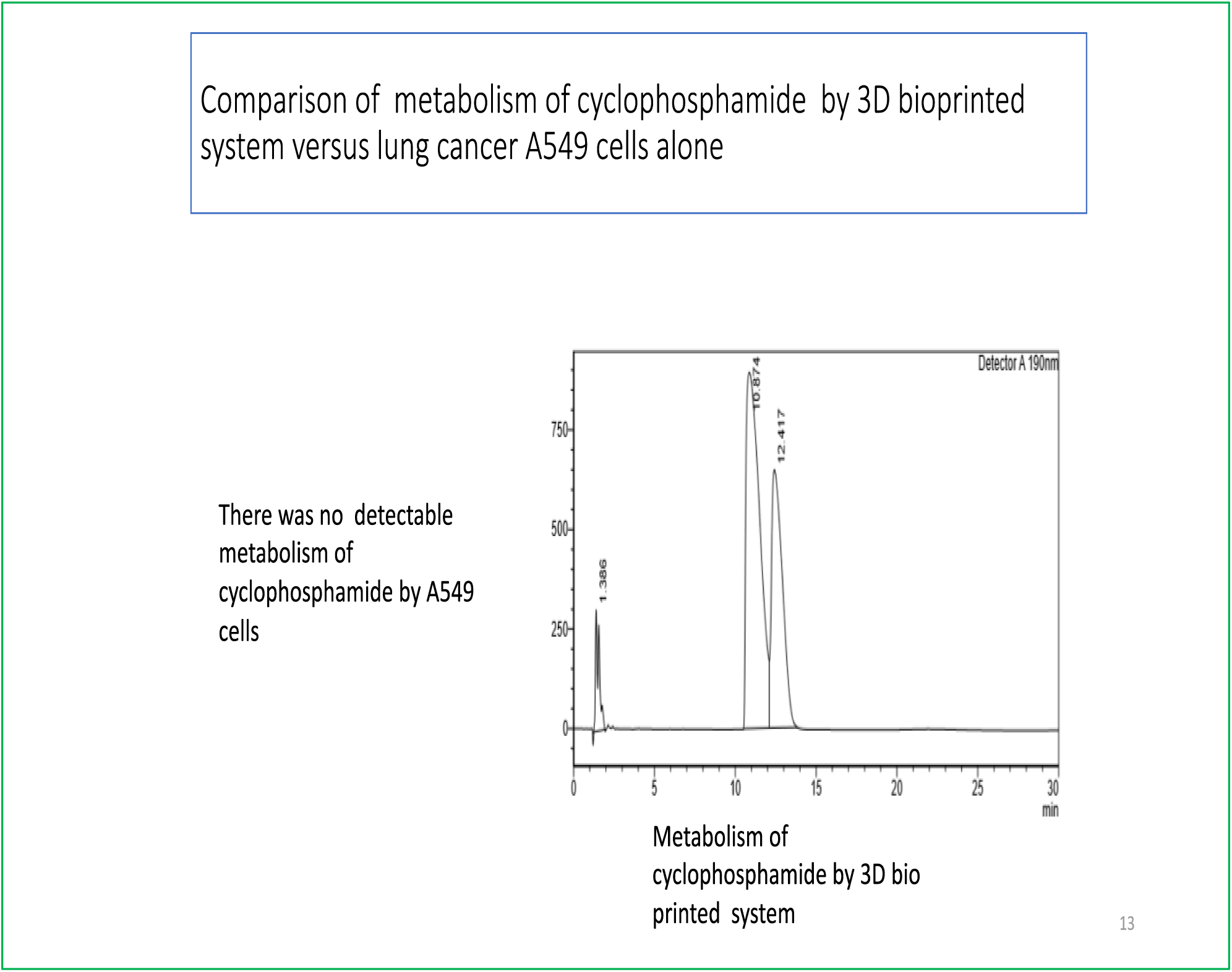

**Figure 7.**
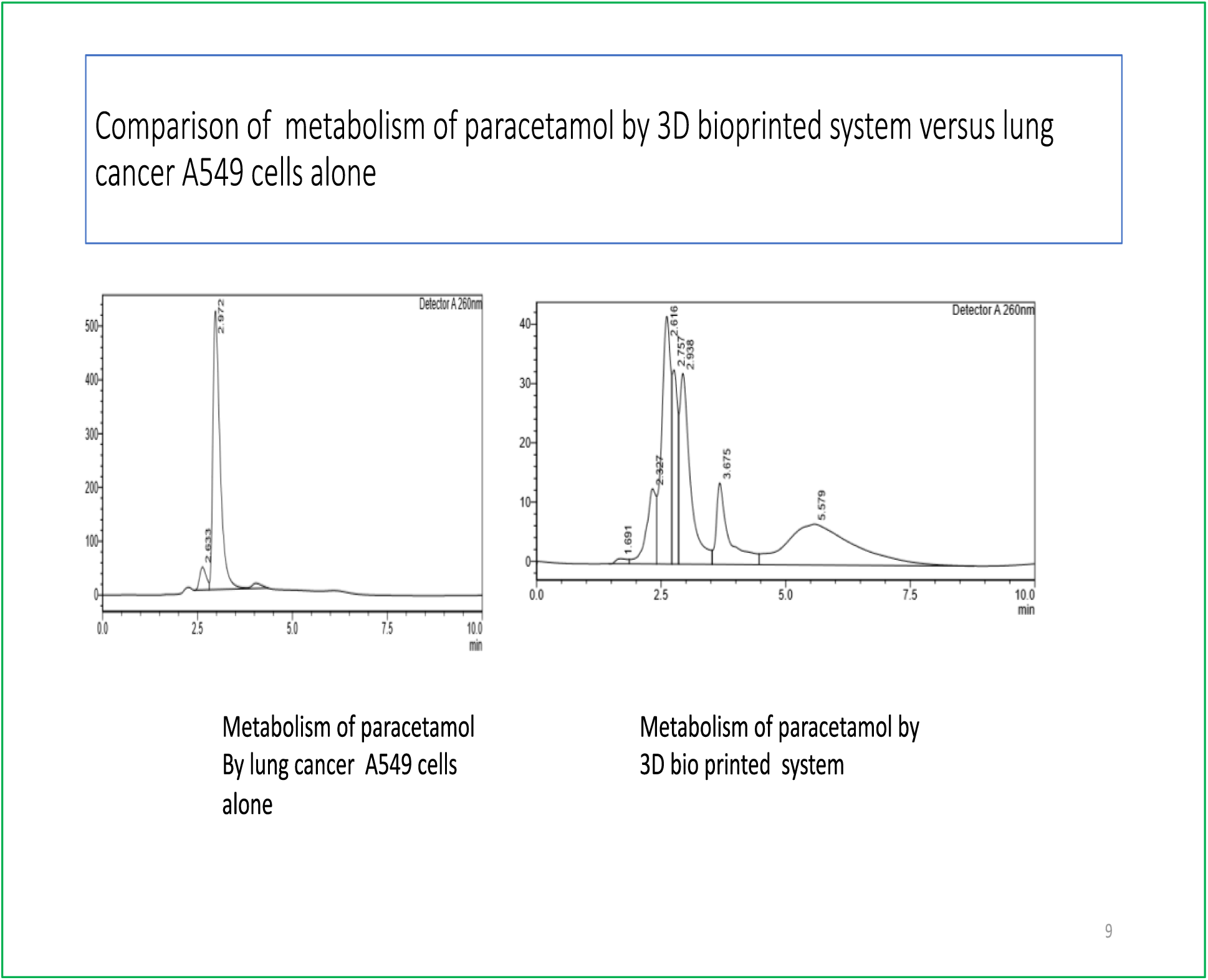

**Figure 8.**
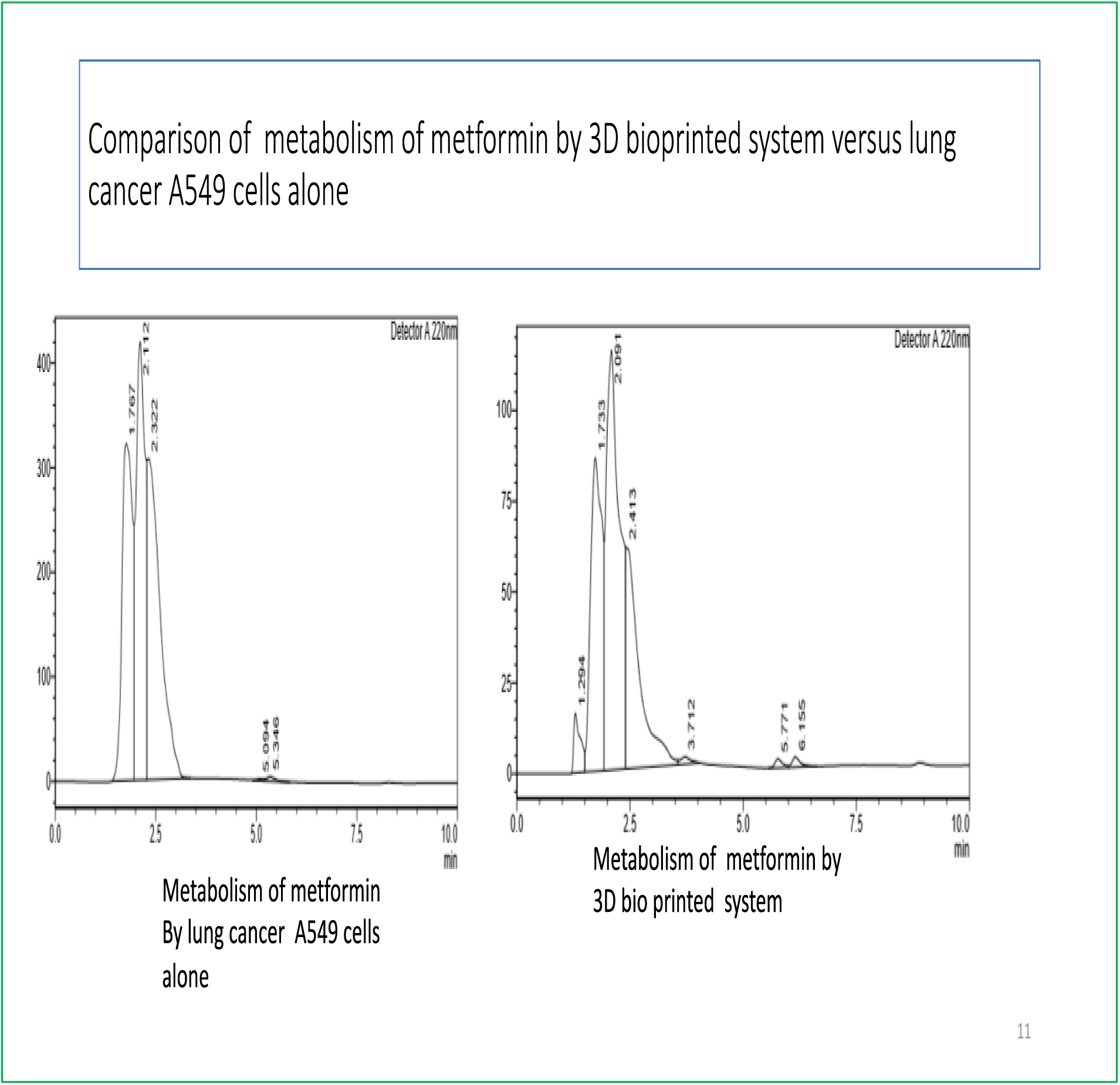

**Figure 5** shows the time dependent metabolism of cisplatin over 6,12 and 24 hours in A549 cells alone versus the 3D bioprinted system. Each metabolite is shown as a bar (orange bar is A549 cells alone and yellow bars represent 3D system). As can be seen there were three major metabolites detected by HPLC. There was one major metabolite and at every time point the amount of this metabolite was higher in 3D system relative to lung cancer cells suggesting superiority of metabolism by the 3D system which contains bioprinted intestinal, liver and lung cells.

In case of the prodrug cyclophosphamide no peaks of metabolites were found in A549 cells but several metabolite peaks were found when incubated with 3D bioprinted system likely because they contain liver cells (**Figure 6**).

Cyclophosphamide is a prodrug which requires metabolic activation by the liver and only the metabolite(s) is active. As expected the lung cells were unable to metabolize this drug and no metabolites were detected. This points out the advantage of profiling drugs in the present 3D bioprinted system versus the traditional way of screening drugs with just the target cells or enzymes.

We also profiled the metabolism of the non-cytotoxic drugs paracetamol, metformin, and indomethacin as shown in **Figures 7-9**. Consistently we found that the metabolite profile generated (compare metabolite peaks generated in both systems) by the 3D system was distinct and superior to that of lung cells alone in terms of the quantity and diversity of metabolites generated. This supports the thesis that an integrated 3D system may be better option to study drug metabolism relative to lung cells (or target cells) alone.

**Figure 9.**
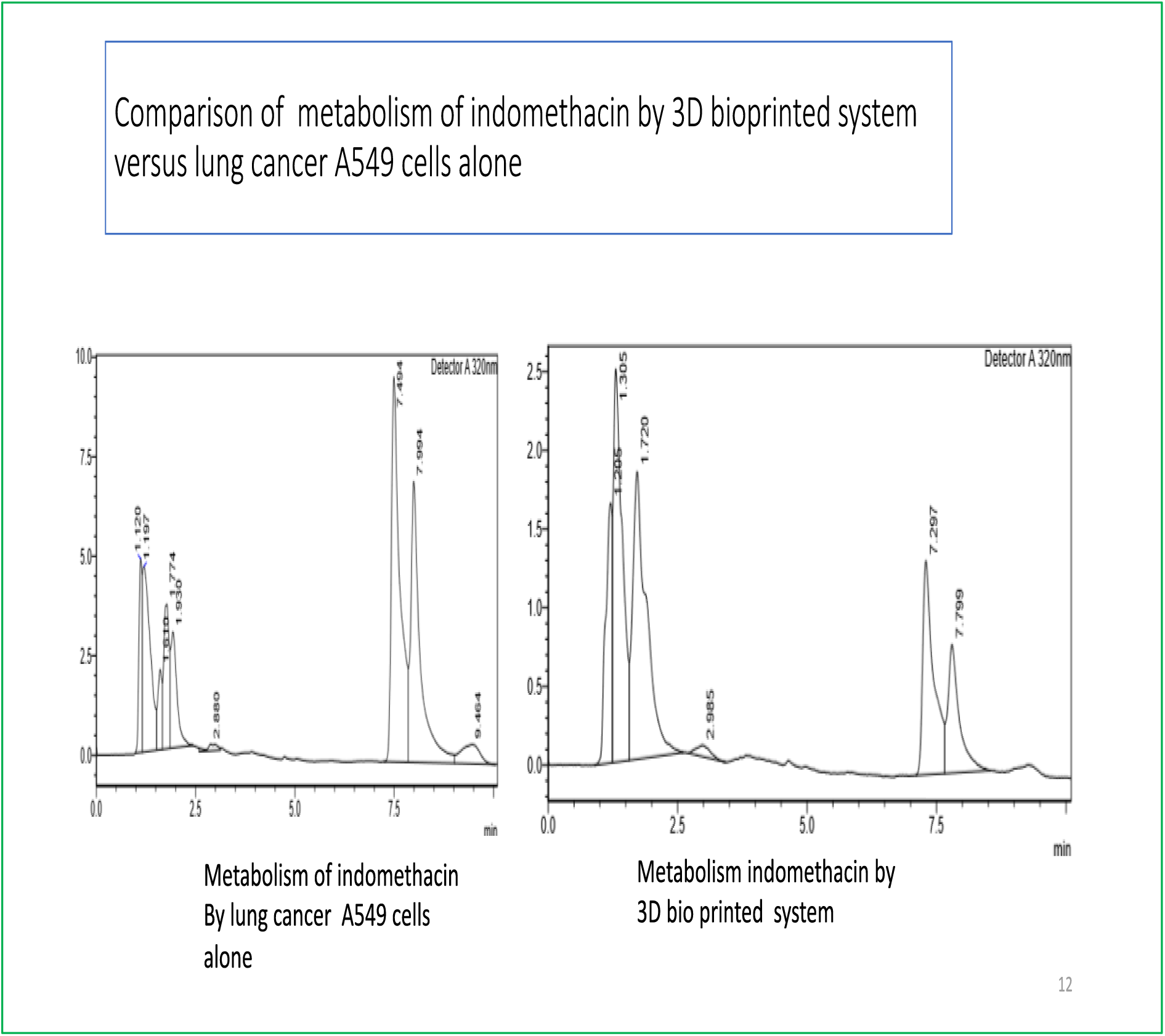

## Discussion

We have developed a 3D bioprinted human cell “disease in a dish” system to study cytotoxic or other drugs against lung cancer. The idea was to create a system that holistically looks at drug efficacy, toxicity and metabolism in a single system rather than the piece meal series of in vitro assays and animal systems.

Our data suggest the feasibility of using this system in identification of lead compounds and repurposed drugs to predict human outcomes. Some of the observations that validate our system are

1. The cytotoxic drugs cisplatin and cyclophosphamide were most active as expected
2. Since they are cytotoxic drugs they were toxic to both target lung cells as well as liver cells as collateral damage, while the non-cytotoxic drugs showed less efficacy or toxicity
3. In terms of drug metabolism, in all cases the metabolites generated by the 3D system were more in quantity and distinct from lung cells alone presumably because of the presence of liver cells in 3D system. A really good example of this was the prodrug cyclophosphamide – lung cells generated no detectable metabolite but the 3D system due to presence of liver cells was able to metabolize the drug

We believe our system can be used to create any human disease simulation system simply by substituting the lung cells by another target cell type. The best use of our system is in the preclinical discovery stage to select right drug candidates to move forward towards preclinical development. This single 3D system could replace the battery of assays that are currently used.

## Acknowledgements

The authors would like acknowledge the assistance of Murali, Rishitha and Tejas in some of the experiments.

## Notes

### Competing Interest Statement

The authors have declared no competing interest.

## Selected References

[1] Application of 3D bioprinting in the prevention and the therapy for human diseases Hee-Gyeong Yi, Hyeonji Kim, Junyoung Kwon, Yeong-Jin Choi, Jinah Jang and Dong-Woo Cho Signal Transduction and Targeted Therapy (2021)6:177 ; https://doi.org/10.1038/s41392-021-00566-8

[2] A high-throughput 3D bioprinted cancer cell migration and invasion model with versatile and broad biological applicability Eric Y. Du, M.A. Kristine Tolentino, Robert H., MoonSun Jung, Joanna N. Skhinas Utama, Martin Engel, Alexander Volkerling, Andrew Sexton, Aidan P. O’Mahony, Julio C. C. Ribeiro, J. Justin Gooding and Maria Kavallaris bioRxiv preprint doi: https://doi.org/10.1101/2021.12.28.474387

[3] Kinetics of Cyclophosphamide Metabolism in Humans, Dogs, Cats, and Mice and Relationship to Cytotoxic Activity and Pharmacokinetics Dominique A. Ramirez, Keagan P. Collins, Allister E. Aradi, Katherine A. Conger, and Daniel L. Gustafson Drug Metab Dispos 47:257–268, March 2019

[4] The anticancer effect of phospho-tyrosol-indomethacin (MPI-621), a novel phosphoderivative of indomethacin: in vitro and in vivo studies Dingying Zhou, Ioannis Papayannis, Gerardo G. Mackenzie, Ninche Alston, Nengtai Ouyang, Liqun Huang, Ting Nie, Chi C. Wong and Basil Rigas Carcinogenesis vol.34 no.4 pp.943–951, 2013

[5] Gleevec (STI-571) inhibits lung cancer cell growth (A549) and potentiates the cisplatin effect in vitro Peilin Zhang, Wei Yi Gao, Steven Turner and Barbara S Ducatman Molecular Cancer 2003, 2:1

[6] Disease modelling in human organoids Madeline A. Lancaster and Meritxell Huch Disease Models & Mechanisms (2019) 12,

